# Inferring light responses of primate retinal ganglion cells using intrinsic electrical signatures

**DOI:** 10.1101/2022.05.29.493858

**Authors:** Moosa Zaidi, Gorish Aggarwal, Nishal P. Shah, Orren Karniol-Tambour, Georges Goetz, Sasi Madugula, Alex R. Gogliettino, Eric G. Wu, Alexandra Kling, Nora Brackbill, Alexander Sher, Alan M. Litke, E.J. Chichilnisky

## Abstract

Reproducing high-acuity vision with an epiretinal implant will likely require inferring the natural light responses of diverse RGC types in the implanted retina, without measuring them directly. Here we demonstrate an approach that exploits intrinsic electrical features of primate RGCs. First, ON-parasol and OFF-parasol RGCs were identified with 95% accuracy using electrical features. Then, the somatic electrical footprint, predicted cell type, and average linear-nonlinear-Poisson model parameters of each cell type were used to infer a light response model for each cell. Across five retinas, these models achieved an average correlation with measured firing rates of 0.49 for white noise visual stimuli and 0.50 for natural scenes stimuli, compared to 0.65 and 0.58 respectively for models fitted to recorded light responses, an upper bound. This finding, and linear decoding of images from predicted RGC activity, suggested that the inference approach may be useful for high-fidelity sight restoration.

## Introduction

Epiretinal implants are designed to provide artificial vision to patients blinded by degenerative photoreceptor disease, by electrically stimulating retinal ganglion cells (RGCs) in a way that transmits useful visual information to the brain.^1,2^ However, the primate RGC population consists of over 20 distinct, intermixed cell types, each of which conveys distinct visual information to distinct targets in the brain.^3,4^ In contrast with this diverse and specific neural circuitry, current epiretinal implants indiscriminately stimulate collections of nearby RGCs without regard to cell type, resulting in unnatural activation patterns (e.g. simultaneous activation ON and OFF cells) and limited vision restoration. No device has been developed that harnesses the accumulated scientific knowledge of the distinct functions of the diverse RGC types to reproduce the neural code of the retina. The development of such a device could improve artificial vision, and could pave the way for leveraging scientific understanding in neural implants of many kinds.

An ideal device that replicates the neural code would first infer, for any given incident visual image, how each RGC would have fired in the healthy retina, and would then electrically stimulate each cell to fire in this predicted pattern. Recently, electrical stimulation of RGCs at single-spike, single-cell resolution has been achieved in the isolated retinas of macaques,^5–7^ and humans,^8^ supporting the possibility of this kind of precise RGC activation. However, inference of the desired RGC activity remains a challenge. While several accurate RGC light response models have been developed (e.g. ^9,10^), these models have always been fitted using measured RGC light responses, which is not possible in the retina of a blind person. Thus, a means of inferring normal RGC light responses in a degenerated retina is needed. One possible approach is to build a model of light response for each cell from intrinsic features of its recorded electrical activity. Indeed, the feasibility of deducing the cell type, a crucial determinant of a cell’s light response properties, from electrical features alone has recently been demonstrated.^11^ However, the reliability of this cell type inference remains a challenge, and the feasibility of inferring a full light response model using only information extracted from intrinsic electrical features has not been tested.

This paper represents the first effort to infer from intrinsic RGC electrical features a full end-to-end quantitative light response model, and to test the results using measured light responses in the same retina. First, a cell type classifier was designed to identify two major RGC types in the macaque retina from electrical features measured on a multi-electrode array, without access to light responses. Then, the electrical footprint of each cell’s action potential on the array was used to infer the receptive field location of each cell. Next, the predicted cell type and predicted receptive field location were combined to infer a full light response model for each cell. Finally, the accuracy of the inferred light response models was assessed by comparing their predictions to the measured light responses of each cell, and also by decoding images from predicted and observed light responses of the full RGC population. The results suggest that the approach produces predictions that largely preserve visually meaningful information and thus may be useful in future high-fidelity retinal implants.

## Results

To infer and test RGC light response models, electrical recordings from isolated macaque retina were performed *ex vivo* using a 512-electrode array system.^12,13^ For each cell, intrinsic features of the recorded electrical activity were used to infer first its type and then a light response model. The accuracy of the inferred model was then assessed by comparing its predictions to observed responses to visual stimuli, and also by reconstruction of visual stimuli from recorded responses.

### Separating ON-parasol and OFF-parasol cells from other cell types

An epiretinal implant targeting certain cell types for stimulation must be able to distinguish them from all other recorded cells. Therefore, the feasibility of separating ON-parasol and OFF-parasol cells, two of the numerically dominant cell types in the primate retina,^14^ from all other recorded types using only intrinsic features of electrical activity was tested. The axon conduction velocity of each cell, estimated from the propagation of spikes along the axon over the multi-electrode array, was effective for this purpose (Fig. 1): parasol cells exhibited a relatively higher axon conduction velocity, while ON-midget, OFF-midget, small bistratified cells, and unidentified cell types, had lower axon conduction velocities,^15,16^ with nearly non-overlapping distributions. The remaining analysis focuses on the ON-parasol and OFF-parasol cells.

**Figure 1:**
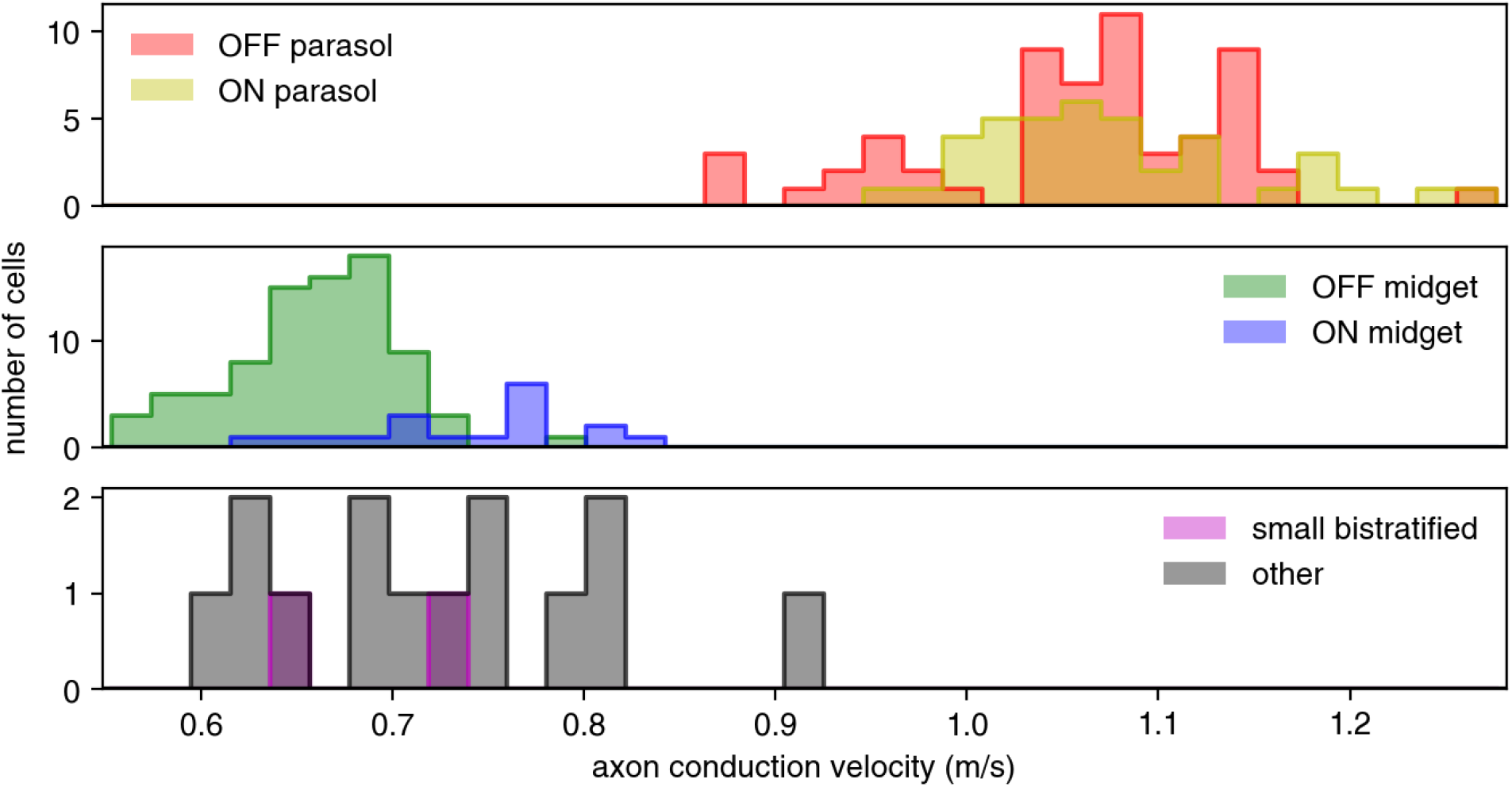
Distinguishing parasol cells by axon conduction velocity. Histograms show axon conduction velocity estimates for 216 of the 946 cells in a single recording, including: 59 OFF-parasol and 40 ON-parasol cells (top); 83 OFF-midget and 18 ON-midget cells (middle); and 2 small bistratified cells and 14 cells of other types (bottom). The remaining 730 cells were excluded on the basis of too few axonal electrodes for an accurate estimate, or a spatial pattern of axonal electrode locations suggestive of a polyaxonal cell.^17^ ON-parasol cells and OFF-parasol cells tend to have higher axon conduction velocities than other cell types.^15,16^

### Cell type classification

Accurately inferring light response models for the ON-parasol and OFF-parasol cells requires separating the cells of the two types using only features of intrinsic electrical activity (i.e., no light response data). Two such features were examined.

The first feature was the autocorrelation function (ACF) of the recorded spikes from each cell. The ACF indicates the firing probability of a cell as a function of time after the occurrence of a previous spike; its form reveals temporal patterns of spiking behavior and tends to be distinct in different RGC types (see Methods).^18,19^ In any given retina, the ACFs of ON-parasol and OFF-parasol cells were usually distinct from one another (Fig. 2A, 2B). When each retina was analyzed individually, k-means (k=2) clustering of the top two principal components of the ACFs resulted in an average 94% *separation* of ON-parasol and OFF-parasol cells (see Methods), with 100% separation achieved for 5 of the 29 recordings. However, due to inter-retina variability in the form of the ACFs, ACFs were less useful for *identification* of the two cell types across retinas (Fig. 2C). For example, cell type classification using machine learning based on ACFs (see Methods) yielded cross-validated classification accuracy of only 65% across 29 retinas. In sum, ACFs alone can be used to separate, but not reliably identify, ON-parasol vs. OFF-parasol cells.

**Figure 2:**
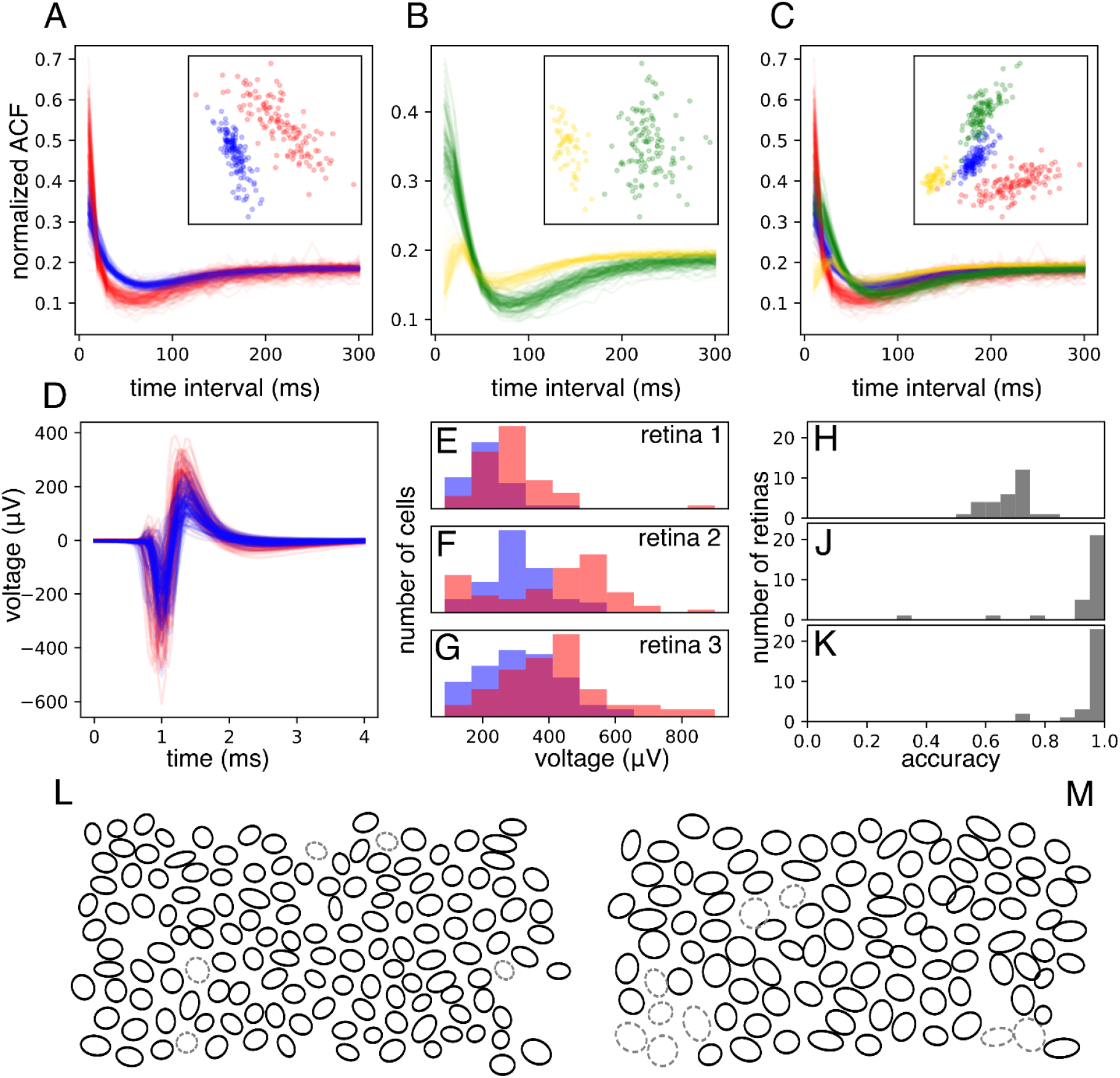
Cell-type Identification among ON and OFF parasol cells. A and B: Comparison of the autocorrelation functions (ACFs) of ON-parasol cells (red in A, yellow in B) and those of OFF-parasol cells (blue in A, green in B), for two retinas (A and B). Insets show projection of each ACF onto the top two principal components across all ON and OFF parasol cells within that retina. C: ACFs from A and B plotted together and projected onto common principal components (inset). D: Main somatic spike waveforms extracted from ON (red) and OFF (blue) parasol cells in one retina. E, F, G: Overlaid histograms of the maximum negative value (the strongest electrode waveform feature, shown here as an example), for ON (red) and OFF (blue) parasol cells within a retina, shown for three separate retinas. H: Histogram of classification accuracy across retinas achieved with spike waveform features alone. J: histogram of classification accuracy achieved using combined approach of ACF-based clustering using the top two principal components and spike waveform based labeling. K: histogram of classification accuracy achieved using an extension of the approach shown in J in which multiple candidate clusterings are considered. L: Mosaic of receptive fields (1SD contours of Gaussian fits) of the OFF-parasol cells of an example retina. Cells incorrectly labeled by the approach shown in K are indicated with gray dashed ovals. M: same as L, but the mosaic of ON-parasol cells for the same retina.

The second feature of intrinsic electrical activity considered for distinguishing cell types was the recorded voltage waveform associated with the action potential, obtained on the electrode recording the largest amplitude spikes from the cell (Fig. 2D). Several features of this spike waveform were examined; among these, the maximum negative voltage of the waveform provided the most accurate segregation of ON-parasol and OFF-parasol cells. Although there was considerable inter-retinal variation in this quantity, ON-parasol cells had consistently higher mean waveform maximum than OFF-parasol cells from the same retina (Fig. 2E, 2F, 2G). The inter-retinal variation was accounted for by normalizing the waveform maxima of all parasol cells in each retina to have zero mean and unit variance. After training on data from 246 recordings from 148 retinas with rapidly generated (imperfect) cell type labels and testing on data from 29 recordings from 29 retinas with carefully curated cell type labels, classification with normalized spike waveform maxima alone achieved a mean accuracy of 63%. Inclusion of spike waveform features recorded from other electrodes did not significantly improve classification accuracy (see Methods). Classification using logistic regression on normalized waveform maximum and 10 additional, similarly normalized features of the waveform (see Methods) increased mean accuracy to 68% (Fig. 2H). Use of neural network based classifiers, applied either to these normalized features or to the waveforms on multiple electrodes, increased accuracy (76%), but not in a way that influenced subsequent inference stages (see below). In sum, features of the spike waveform alone imperfectly identify ON-parasol vs. OFF-parasol cells.

These results suggest a combined approach to cell type identification: separate the ON-parasol and OFF-parasol cells of each retina using their ACFs, and then identify the clusters using features of the spike waveform. Using k-means clustering on the top two principal components of the ACF and then identifying clusters by “voting” using the spike waveform classifier predictions achieved 80% mean (93% median) accuracy on the training retinas and 93% mean (97% median) accuracy across the 29 test retinas (Fig. 2J). In sum, a combination of ACF and spike waveform information can reliably identify ON-parasol and OFF-parasol cells.

To further improve the accuracy of cell type classification, ACF based clustering was optimized. While k-means clustering on the first two principal components (PCs) of ACFs was more accurate than any other fixed number of PCs, using a different number of top PCs for different retinas was more accurate. Furthermore, despite inter-retina variability, the clustering accuracy of some retinas improved with including as clustering features projections onto the top PCs of the pooled ACFs of all retinas, termed *global* ACF PCs, in addition to projections onto the top PCs from the same retina, termed *local* ACF PCs. Therefore, in the optimized approach, varying numbers of top local ACF PCs and global ACF PCs were used to generate 100 candidate clusterings per retina. Without knowledge of true cell types, the optimal choice among the candidate clusterings was taken to be the clustering that maximized agreement with the initial spike waveform classifier predictions while avoiding excessive imbalance in the predicted ratio of cell types (see Methods). With selected candidate clusters labeled according to spike waveform classifier votes as before, this optimized approach increased overall cell type prediction accuracy to 83% mean (95% median) accuracy on the rapidly labeled training data and to 95% mean (97% median) accuracy on the carefully labeled testing data (Fig. 2K).

Substituting a neural network based classifier utilizing the waveforms of multiple electrodes did not increase this final accuracy: 79% mean (94% median) accuracy on training data and 93% mean (98% median) accuracy on test data.

### Inferring a light response model for each cell

To infer the light response properties of each RGC, the inferred cell type was used to construct a linear-nonlinear Poisson (LNP) cascade light response model, and the accuracy of model predictions was assessed.^20^ In the LNP model, the visual stimulus passes first through a linear spatiotemporal filter, which integrates across nearby space and recent time to produce a univariate generator signal over time. This generator signal passes through a scalar nonlinear function to determine the time-varying firing probability, and spikes are generated from this probability according to an inhomogeneous Poisson process.

To estimate the model parameters for each cell using only its inferred type and intrinsic features of its electrical activity, a two-step procedure was used. First, the nonlinearity function of each RGC was assumed to be the average nonlinearity function for all cells of its inferred type within that retina. Second, the linear spatiotemporal filter of each RGC was assumed to have the average shape for its inferred type, but translated in space to an inferred receptive field center location. To infer the receptive field center of a cell, first its soma center was inferred from its electrical image (EI), the spatiotemporal electrical footprint of the action potential on the array. The soma center was computed as the weighted average of the locations of the maximum spike amplitude electrode and its neighboring electrodes, with weight equal to the spike amplitude of that electrode. Exploiting the fact that the receptive field of a cell tends to lie over its soma,^21,22^ soma locations were then mapped to visual input space to obtain estimates of receptive field centers (Fig. 3; Fig. 4 center). Taken together, these inferred model parameters constituted an inferred LNP model for each RGC.

**Figure 3:**
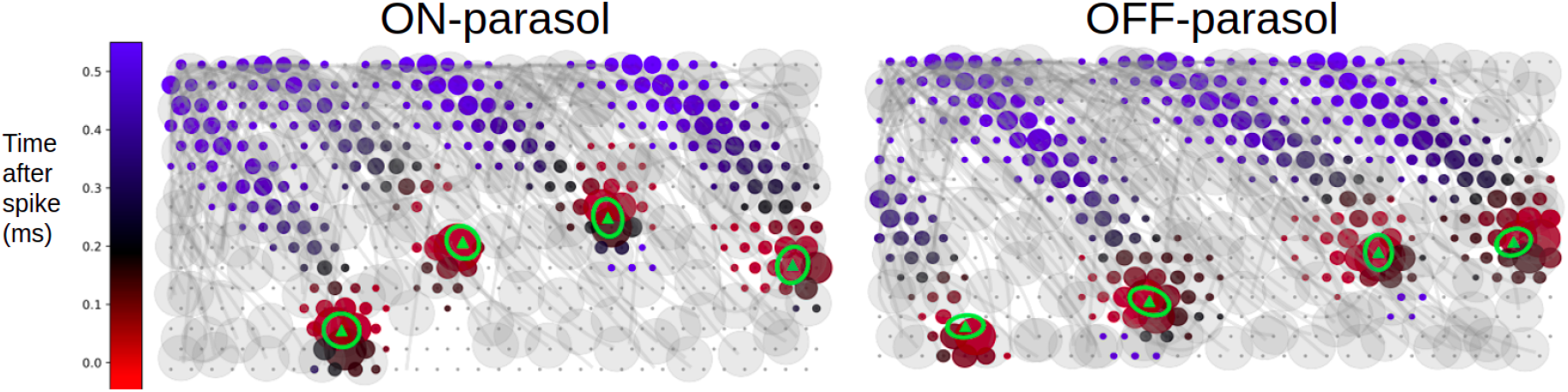
Inferring receptive field centers from electrical images (EIs). Example EIs with color indicating the timing of propagation of the spike from the soma down the axon, and circle size indicating the spike amplitude, at each electrode. Receptive fields obtained using light responses (green ellipses with triangle centers) mapped into the plane of the electrode array demonstrate the tendency of receptive fields to lie over the soma.

**Figure 4:**
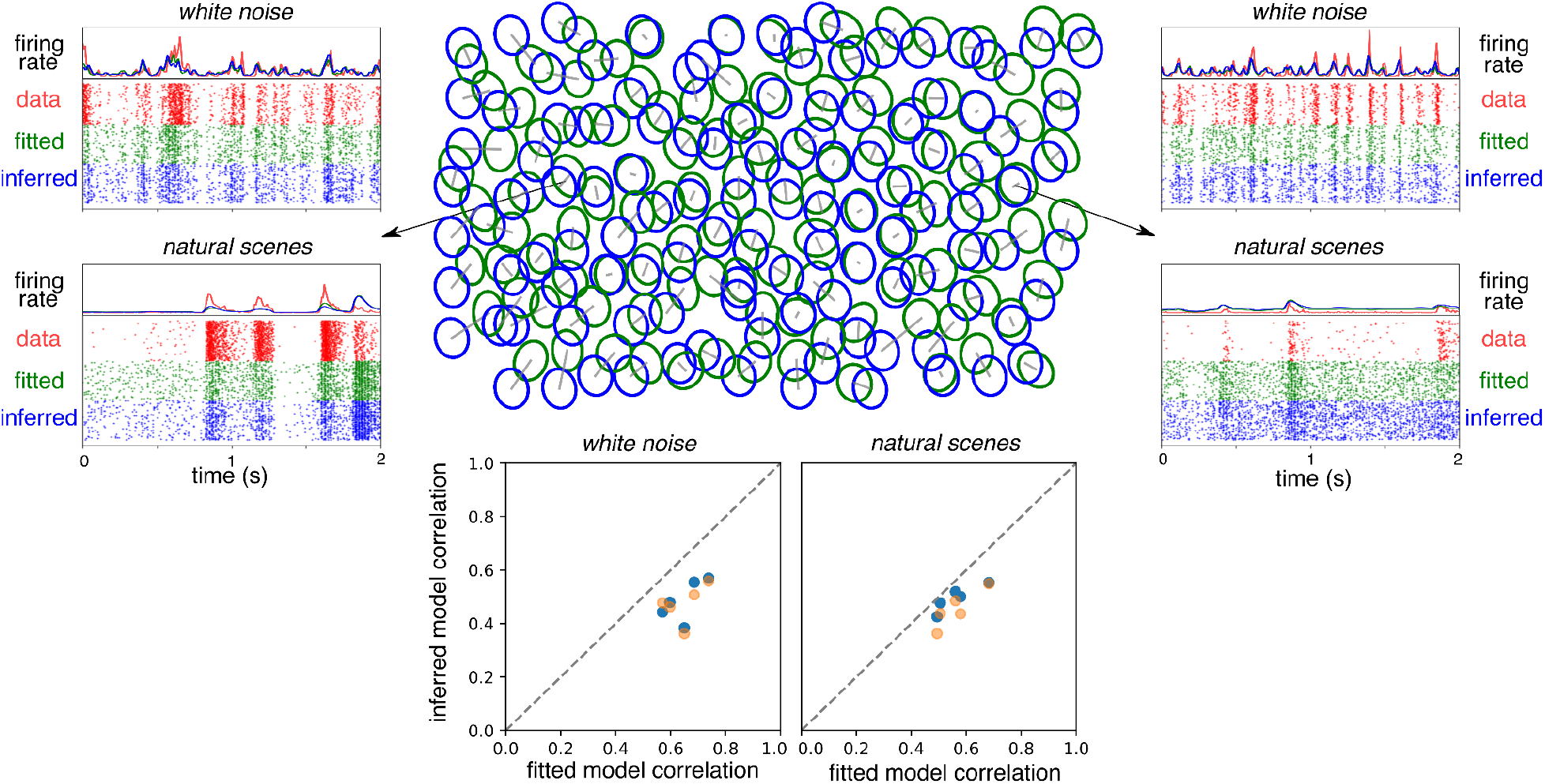
Encoding model inference. Top center: mosaic of true receptive fields (RFs), obtained by fitting to responses to white noise visual stimuli (fitted, green), overlaid with the mosaic of receptive fields inferred from electrical features (inferred, blue), for the OFF-parasol cells in one retina. Each receptive field is shown as 1 SD contour of the Gaussian fit to the corresponding linear spatial filter. For each cell, a gray bar connects its true and inferred receptive field center. Top left and Top right: sample firing rate and spike raster across trials for observed responses (data), fitted model predictions (fitted), and inferred model predictions (inferred) for two example cells. Bottom: average performance (correlation between predicted and observed firing rates) across parasol cells for white noise (left) and natural scene stimuli (right) for five retinas, using either the retina-specific cell type averages of RF shape, time course, and response nonlinearity for model inference (blue), or the across-retinas cell type averages (orange)

### Testing light response models

To assess the accuracy of the inferred light response model for each RGC, the response to a visual stimulus predicted by the inferred model was compared to a measured response to that stimulus. Both natural scenes and white noise stimuli were tested. To control for the stochasticity inherent in RGC responses, each visual stimulus was repeated 30 to 60 times. The inferred and true responses were similar for most cells (Fig. 4), with an average Pearson correlation coefficient of 0.50 for white noise stimuli and 0.53 for natural scenes stimuli across five retinas (Fig. 4, bottom, blue).

To distinguish errors in model predictions attributable to imperfectly inferring model parameters from errors attributable to limitations of the LNP model itself, for each RGC, observed light responses were also compared to the predictions of a model that was fitted to held-out data.

Across the same five retinas as above, the true firing rate and fitted model prediction firing rate had an average Pearson correlation coefficient of 0.65 and 0.58, for white noise and natural scenes stimuli respectively (Fig. 4, bottom). These fitted model predictions establish an upper bound on the accuracy that can be achieved by inferring the LNP model as above.

To test the effect of inter-retina variability on prediction of light responses, the above analysis was repeated with the average spatiotemporal filter shape and nonlinearity per cell type averaged across multiple retinas, rather than across cells within the retina being tested. Across the same five retinas, the models produced a mean correlation between inferred and true responses of 0.49 for white noise stimuli and 0.50 for natural scene stimuli (Fig. 4, bottom, orange). Overall, these results indicate that cell type is highly predictive of light response properties, but that retina-specific information is also significant,^23^ a potential challenge for the development of high-fidelity implants (see Discussion).

### Testing for preservation of visual information

To test further how effectively the inference of light response models captures visual signaling by the retina, linear decoding of simulated natural scene stimuli was performed, using the RGC activity predicted by both fitted and inferred encoding models. The linear decoder can be interpreted as a simple model of downstream visual processing by the brain, which must make inferences about the visual scene to drive behavior (see Discussion). The stimuli consisted of natural images (ImageNet database), presented for 100 ms, interleaved with a uniform gray background with intensity equal to the mean across images. The interval between image presentations was 400 ms, to ensure independence of the evoked responses to each image (see Methods). A collection of images and responses was designated as training data for the decoder, and other images and responses were designated as test data. The training data were used to learn a linear transformation that decoded the images from the combined responses of all recorded ON-parasol and OFF-parasol cells, with least squared error (see Methods and Brackbill et al.^24^). This linear decoder was then applied to the test responses, and the resulting decoded images were compared to the original images. The similarity between each original image and its decoded version was measured by the correlation coefficient. Next, the light response model inference method presented above was used to infer RGC responses to the images, and the same linear decoder was applied to inferred model predictions.

Across 150 test images, the mean correlation between true images and images decoded from the fitted model RGC responses was 0.81, and the mean correlation between the true images and the images decoded from inferred RGC responses was 0.55. This 0.55 mean correlation increased to 0.69 if the true rather than inferred cell types were used in inferring RGC responses. For comparison, the correlations obtained with randomly permuted true images was <0.001. Thus, RGC response inference captures most of the visual information captured by a best fit encoding model, an upper bound (Fig. 5).

**Figure 5:**
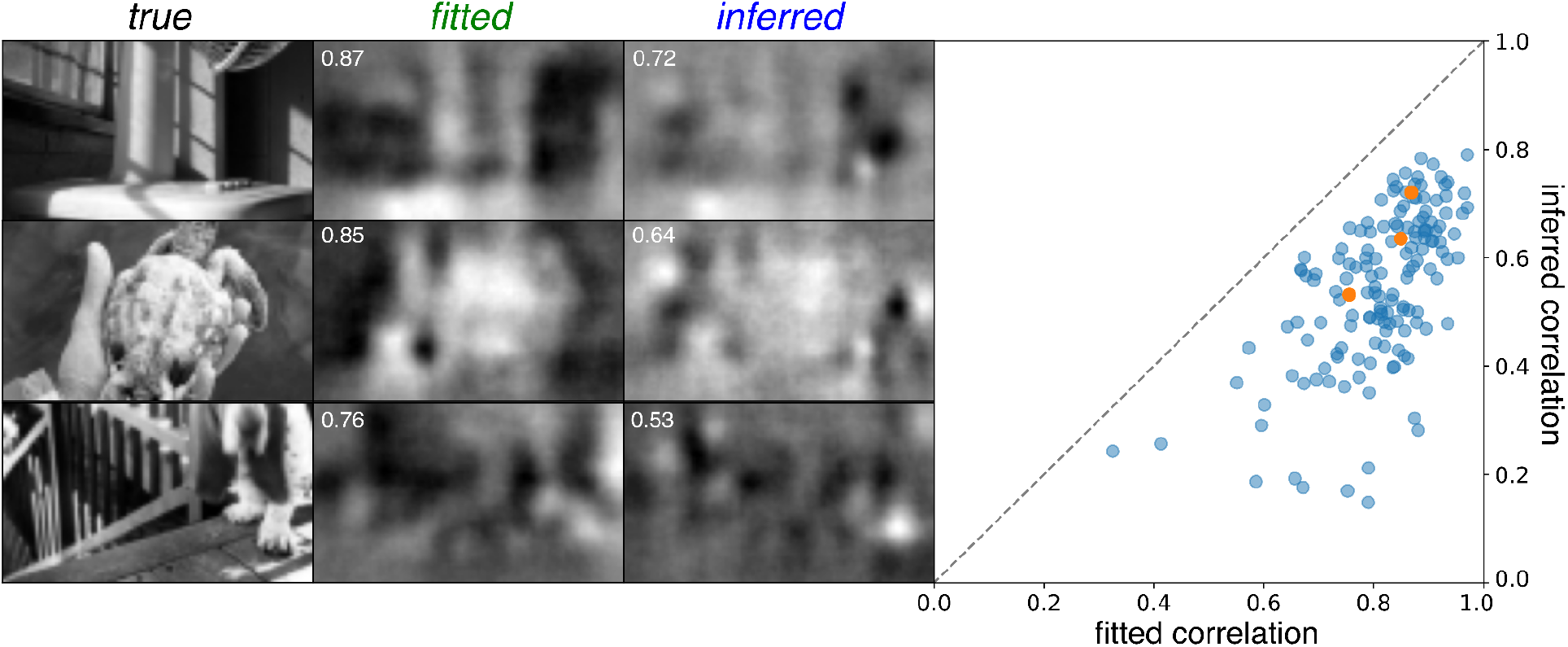
Linear decoding of responses. Left: three example reconstructions from fitted model (fitted) and inferred model (inferred) responses compared to the test images used to elicit the responses (true). Performance scores (correlation with true image) is provided for each reconstruction (white inset text). Right: scatterplot of performance scores of inferred model reconstructions compared to fitted model reconstructions across 150 test images, with the three example images indicated (orange).

## Discussion

A two-step approach was used to infer RGC light response models using only intrinsic features of electrical activity: first, electrical features were used to infer cell types, and second, cell type information was combined with additional information from electrical features to infer a light response model for each cell. Although the first step, cell type inference, has been attempted previously,^11^ the 95% accuracy on ON-parasol versus OFF-parasol cell discrimination achieved here using a new and more interpretable classification approach and much larger data set represent a substantial improvement. The second step, light response model inference, has not been attempted previously. The predictions made by the inferred light response models closely matched the predictions of models fitted to measured light responses, validating the approach. Decoding of stimuli from responses further indicated that the inferred models tend to capture the essential properties of normal visual signaling by the retina. More generally, these findings indicate that there is enough information present in the intrinsic electrical features of a neuron to understand its cell type and detailed function within a circuit, a conclusion with potential implications for many brain-machine interfaces.

In the retina, the findings support the feasibility of a new generation of high-fidelity implants that replicate the neural code. Such devices would first use recorded spontaneous activity to infer the RGC firing that would have occurred in the intact retina, and then would precisely evoke that normal firing pattern using electrical stimulation. Nevertheless, there are several caveats, both for light response inference and for its application to the design of retinal implants.

### Caveats to inferring light responses from electrical features

The present approach is limited by inter-retina variability in RGC encoding properties. The analysis focused first on the inference problem in a single retina, by using the average spatial filter, temporal filter, and nonlinearity function of cells of that type in the same retina in order to model light responses. However, this is unrealistic in the clinical setting. A more realistic approach – using these features averaged across multiple retinas – resulted in less accurate model inference. However, recent work suggests a third approach. Specifically, it may be possible to infer retina-specific encoding properties using psychophysical measurements in the implant recipient. In these measurements, the visual percepts generated by electrical stimulation designed to emulate different light response models could be compared.^23^ Notably, the proposed psychophysical approach also requires cell type classification, highlighting the importance of the cell type inference developed here.

Most of the present analysis was limited to ON-parasol and OFF-parasol cells, two of the numerically dominant cell types in the primate retina. A more complete approach will require expanding cell type classification and light response model inference to other cell types. In particular, ON and OFF midget cells will be important, given their high density^25^ and consequent importance for high-acuity vision. Identifying and targeting additional cell types may be more challenging.

Errors in identifying the receptive field location of each cell from its electrical image contributed substantially to errors in light response inference (not shown). Therefore, future work will prioritize techniques to more accurately infer the receptive field location. For example, inferring the dendritic field extent of a cell from its EI, in addition to inferring the somatic center, may improve inference.^22^

The current approach is also limited by the assumption that cells of the same type all have the same light response properties, except for receptive field location. Future work will explore whether light response model inference can be more tailored to individual cells. For example, the electrical image of a cell may contain cell-specific information about the shape or orientation of the receptive field. This may be particularly important for cell types with receptive fields that are not smooth and circularly symmetric.^19^

Finally, the present analysis is limited by the exclusive use of the linear-nonlinear-Poisson model of light responses. The choice of encoding model places an upper bound on the accuracy of inferred model predictions, and the performance of LNP models fitted to measured responses was thus included for comparison. More complex models, such as those that incorporate the temporal structure of spike trains,^9^ cross-correlations in cell firing,^10^ or nonlinear subunit structure in receptive fields,^26^ could potentially offer higher performance,^27–29^ but inferring the parameters of such models would require developing new methods that do not require access to light responses.

### Caveats for application to a future bi-directional retinal implant

One caveat for applying this approach to the design of a retinal implant is that visually-driven electrical activity was used to measure electrical features: in a blind retina, this would have to be accomplished with spontaneous firing, which occurs at a lower rate^30^ and could produce slightly different electrical features. Specifically, differences in the ACFs or EIs could complicate cell type classification or model inference. The higher firing rate observed during visual stimulation indicates that the ACFs must differ to some degree, but further work is needed to fully characterize these differences. A smaller difference is expected between the EIs obtained from spontaneous and visually-driven activity, since the action potential of a cell is generally highly stereotyped. Nevertheless, small variations in recorded EIs with varying visual stimuli have been observed in certain RGC types in macaque^19^ and other species.^31,32^

In addition, the electrical features of spontaneous recordings in the healthy retina may not accurately represent the electrical features of cells in the degenerated retina, the target for retinal implants. Fortunately, the present approach to cell type classification only requires that the ACFs of different cell types in a given retina be distinct from each other; the exact nature of the ACFs does not affect inference because the EIs provide the final step of cell type identification. Interestingly, in a rat model of retinitis pigmentosa, the ACFs of ON and OFF cells become more distinct during retinal degeneration (unpublished data). On the other hand, the present cell type classification approach does depend on the exact nature of EIs relative to the EIs on which the classifier has been trained. Since the EI of a RGC largely reflects its intrinsic properties, such as location and density of ion channels, it may be relatively preserved in diseases of photoreceptor degeneration, although there is some evidence to suggest otherwise.^33–35^

An additional caveat to applying these findings to a future human implant is that they were obtained in macaque retina. Promising recent results obtained from human retina recordings *ex vivo*, and comparison to hundreds of macaque *ex vivo* recordings made in a similar experimental setting, reveal remarkable functional similarity between human and macaque retina, including in responses to light^23,36,37^ and to electrical stimulation.^8^

Finally, realization of the envisioned high-fidelity retinal implant will require development of novel hardware that can perform high-density, large-scale recording and stimulation, while maintaining the electrodes in close apposition with the retina, and communicating with an external controller, all in an encapsulated device that operates within a strict power budget to prevent overheating the retina. No present-day hardware can exploit the analytical methods presented here, but efforts are underway to develop such a device.^38,39^

### Future work

One exciting possibility for improving cell type classification in a future implant is to employ psychophysical measurements. Psychophysical methods have previously been used to understand phosphenes elicited by axonal and somatic stimulation with the coarse electrode arrays of present-day epiretinal implants.^40^ For cell type classification, an individual cell could be electrically stimulated using a high-fidelity implant, and the implanted person could report aspects of their perception to help determine the cell type. Alternatively, a group of unidentified cells that are believed to be of the same type based on ACF clustering could be simultaneously stimulated, generating a stronger visual percept for classification. This ACF-based clustering and psychophysics-based cluster labeling approach would be particularly useful if EIs change during degeneration but ACFs of different cell types remain distinct (see above).

The retina is an ideal first application of the present approach, because its neural code is well characterized and its accessibility permits electrical stimulation at cellular resolution. Even so, the overall approach -- inferring the natural function of neurons from intrinsic electrical features, and then reproducing their natural function through electrical stimulation -- could be applicable to other parts of the nervous system. For example, recent work^41^ used the spike waveforms of cortical neurons to reveal greater cell type diversity than had previously been appreciated,^42–44^ a finding that could lead to more precise cell type targeting in cortical implants. These and related approaches could contribute to a new generation of high-performance neural interfaces.

## Acknowledgements

We thank J. Carmena, K. Bankiewicz, T. Moore, W. Newsome, M. Taffe, T. Albright, E. Callaway, H. Fox, R. Krauzlis, S. Morairty, and the California National Primate Research Center for access to macaque retinas. We thank Kristy Berry, K. Williams, B. Morsey, J. Frohlich, and M. Kitano for accumulating and providing macaque retina metadata. We thank R. Vilkhu, P. Vasireddy, M. Hays, A.J. Phillips, S. Cooler, F. Rieke and the entire Stanford Artificial Retina team for helpful discussions. We thank Stanford Medical Scholars Fellowship Program (MZ), Pew Charitable Trusts (AS), Google internship and Student Researcher programs (NPS), Research to Prevent Blindness Stein Innovation Award, Wu Tsai Neurosciences Institute Big Ideas, NIH NEI R01-EY021271, NIH NEI R01-EY029247, and NIH NEI P30-EY019005 (EJC), NIH NEI F30-EY030776-03 (SM), NSF IGERT Grant 0801700 (NB) and NIH NEI F31EY027166 and NSF GRFP DGE-114747 (NB) for funding this work.

## Methods

### Experimental overview

Multielectrode array recordings from isolated macaque retina were obtained using previously-described procedures.^6,45–48^ Briefly, eyes were taken from macaques terminally anesthetized by other laboratories in the course of their experiments. The extracted eyes were hemisected and the vitreous removed in room lighting. The posterior portion of the eye was then kept in darkness in oxygenated Ames’ solution (Sigma) at 33°C. Small (∼3 mm) segments of retina were then isolated under infrared illumination and mounted on a custom multielectrode array, RGC side down. The array consisted of 512 electrodes, 8-10 μm diameter, arranged in an isosceles triangular lattice with 60 μm spacing between electrodes in each row and between rows. During a recording, voltages were simultaneously recorded from all 512 electrodes, bandpass filtered (43-5000 Hz), and digitized (20 kHz). During recording the retina was superfused with oxygenated Ames’ solution at 33-35°C.

### Visual stimulation and ground truth cell type determination

During recordings, visual stimuli were presented on a computer display refreshing at 120 Hz, as described previously.^24^ The mean image intensity was low photopic (∼500-1000 photoisomerizations/photoreceptor/sec). The image was focused on the retina using a microscope objective. White noise visual stimuli consisted of a grid of pixels, each 44×44 or 88×88 μm at the retina, updating at 30 or 60 Hz. At each spatial location, the intensity of each monitor primary was selected from a binary distribution (minimum or maximum intensity) over time, and these were either yoked (producing black-and-white spatial noise patterns) or independent (producing colored spatial noise patterns). Analysis of spatial, chromatic, and temporal light response properties of RGCs with these stimuli were used to determine the ground truth cell types, as previously described.^13,19,20,49^ Natural scenes visual stimuli consisted of images from the ImageNet database^50^ converted to grayscale and downsampled to either 320×160 pixels, with each pixel 11×11 μm on the retina, or to 160×80 pixels, with each pixel 22×22 μm on the retina. For assessment of preservation of visually significant information (Fig. 5), each natural scenes image was shown for 100 ms, followed by a uniform gray display at mean image intensity for 400 ms, for a total of 500 ms per image.^24^ For direct comparison of predicted and observed responses (Fig. 4), natural scenes images were shown for varying duration, without interposing gray display, and with intermittent small translations intended to simulate eye movements.^24^

### Extraction of autocorrelation functions and electrical images

Spikes from distinct cells were identified and separated using techniques previously described.^12^ The spiking behavior of each RGC under white noise stimulation was summarized by its autocorrelation function (ACF). The ACF of a cell was computed by counting the total number of spike pairs separated by a given time interval, for each of 30 evenly spaced intervals between 0-300 ms. This vector of spike counts was then normalized (L2). This normalization accentuated cell type differences in the ACFs. The spikes from each cell were also used to compute its *electrical image* (EI). The EI consists of the average voltage waveform recorded on each electrode during the spikes of a given cell.^12^

### Distinguishing parasol cells from other cells by axon conduction velocity

Axon conduction velocity was used to demonstrate the feasibility of distinguishing ON and OFF parasol cells from all other recorded cell types without access to light responses.^8,15^ To estimate the axon conduction velocity of a cell, the time of maximum negative voltage within the EI waveform was determined for each electrode, after upsampling tenfold in time. Then, for all pairs of electrodes recording an axonal signal, a conduction velocity estimate was computed by dividing the distance between the electrodes by the time difference. Axonal electrode pairs with negative peaks less than one original sample apart (0.05 ms) were excluded. The overall conduction velocity for the cell was computed as a weighted average of the conduction velocity estimates of all axonal electrode pairs, with the weighting given by the product of maximum amplitudes. The top ten weighted pairs were used in the average, and cells with fewer than six axonal electrodes were excluded from analysis. Cells with axonal electrodes more than 90 degrees apart relative to the soma center were also excluded from axon conduction velocity estimation because they likely were polyaxonal amacrine cells.^17^ Subsequent analysis focused only on ON-parasol and OFF-parasol cells.

### Classification of parasol cells as ON or OFF using electrical features

The first step in classifying ON-parasol vs. OFF-parasol cells in each recording using only their electrical features was to assign a preliminary label to each using a classifier operating on the EI waveform on the maximum amplitude electrode. The classifier used logistic regression applied to selected features of the main spike waveform. To extract these features, first the main spike waveform was upsampled in time by a factor of ten to obtain a waveform *f*(*t*), where *f*(*t*) is the voltage at time *t*. Given that somatic waveforms tend to consist of a negative deflection followed by a positive deflection, temporal information was summarized by six time points: *t*_*n*_, time of minimum voltage, *t*_*p*_ the time of maximum voltage, *t*_*n,left*_ first time point before *t*_*n*_ with voltage amplitude less than half that at *t*, _*n*_ *t* _*n,right*_ first time point after *t* _*n*_ with voltage amplitude less than half that at *t* _*n*_, and analogously defined *t*_*p,left*_ and *t* _*p,right*_. Then, using these six time points, the 11 hand-crafted features were then computed as follows: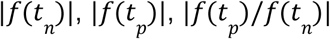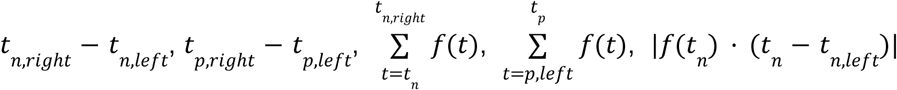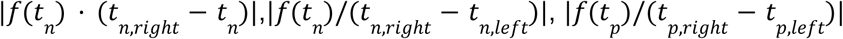
To reduce the effect of inter-retinal variability, values for each feature were normalized to zero mean and unit variance across all the parasol cells within each retina. Feature weights were learned by training on 246 recordings from 148 retinas with rapidly generated (imperfect) cell type labels, and the classifier was tested on 29 recordings from 29 retinas with carefully curated cell type labels. Inclusion of the waveform on apparent dendritic electrodes was also considered as a potential feature but was found to not significantly improve performance (69% accuracy with these additional features and 68% without), therefore only the 11 handcrafted features of the main spike waveform described above were used as features for EI-based classification.

The moderately high accuracy obtained by EI-based classification was significantly improved by using the predictions of EI-based classification to label ACF based clusterings. Two approaches were considered. In the simpler approach, cells of each recording were clustered using k-means (k=2) clustering on projections onto the top two principal components of ACFs for each recording. The clusters were then labeled in the way that maximized agreement with the preliminary labels. In the second approach, projections of ACF onto a variable number of top principal components (1 to 10) of the ACFs of that recording along with a variable number of top principal components (1 to 10) of ACFs pooled across recordings were considered as possible clustering features. K-means (k=2) was performed on each possible combination to generate a total of 100 candidate clusters per recording. Each candidate cluster was assigned cell type labels in the way that maximized agreement with the preliminary labels as with the first approach. The final clustering was chosen among the candidate clusterings, without knowledge of the true labels, as the candidate clustering that maximized agreement with the preliminary labels and that did not predict a cell type imbalance of greater than 0.05 and 0.95. This cutoff value (5%) and the relative weight of the penalty for violating this cutoff value were learned as hyperparameters from the training data.

### Inferring receptive field centers from electrical images

To aid in inferring light response models from electrical features, receptive field centers were inferred from EIs. The soma center location of each cell was taken to be at the electrode with the most negative voltage in the EI. Assuming that the receptive field of a cell tends to lie over its soma,^51,52^ these soma centers were then mapped onto visual stimulus space to estimate the receptive field centers. Such a mapping was necessary because the soma centers were defined in terms of a physical location on the electrode array, whereas receptive field center locations were defined in the coordinate space of the visual stimulus. The mapping from electrode space to visual stimulus space was the least squares affine mapping from the inferred soma centers to the true receptive field centers. In principle, this mapping could have been estimated using the measured relationship between the visual stimulus image and retina. In practice, precise estimation of the mapping using this approach is difficult, particularly due to slight deformations of the photoreceptor layer when the retina is pressed against the electrode array. Determining the mapping from electrode space to visual stimulus space without access to the true receptive field centers presents a challenge for a future retinal implant. However, a simple constant offset in the receptive field centers would likely be more tolerable in the setting of an implant.

### Inferring an encoding model

The cell type and the inferred receptive field center were combined to infer a Linear-Nonlinear-Poisson (LNP) light response model for each parasol cell in five test retinas.^20^ The linear portion of the model was decomposed into separate spatial and temporal components. The overall model thus consisted of the following sequential components: a linear spatial filter, a linear temporal filter, a scalar nonlinearity, and an inhomogeneous Poisson process with an instantaneous spike rate given by the output of the nonlinearity. The nonlinearity was of the form *f*(*x*) = *a* * *N*(*bx* - *c*), where a, b, and c were parameters of the model, and *N* is the cumulative normal function. To infer the model parameters for a cell, the cell was assumed to have the average linear spatial filter shape, temporal filter, and nonlinearity for its cell type.

Both the retina-specific cell type averages and the cell type averages across five retinas were considered for these components. The ground truth model parameters used for these averages were the parameters obtained by fitting the model to the white noise light responses of each cell independently as previously described.^20^ The spatial filter was centered on the inferred receptive field center.

To test the accuracy of the inferred encoding models, model predictions were compared to the observed responses for both white noise and natural scenes visual stimuli. The visual stimuli were of the form described above. Each test visual stimulus was repeated 30 to 60 times to capture stochasticity in firing across trials. The similarity between the true and predicted responses was assessed using the correlation coefficient between the average true and predicted firing rate across trials. For comparison, the similarity between the true response and the predictions of an encoding model fitted to a separate white noise visual stimulus was assessed in the same way.

### Natural scenes decoding

To assess the effectiveness of inferred models for conveying visual information, linear decoding from fitted model-predicted and inferred model-predicted natural scenes light responses was performed, utilizing previously described methods^24^ as summarized below.

First, a linear decoder was trained using the simulated light responses to 10,000 natural images as predicted by the white noise fitted encoding model of a retina. The predicted light response to each image was summarized by the expected number of spikes within 150 ms of when the image was displayed, for each parasol cell. The linear decoding mapped the expected spike counts from all cells to each pixel of the visual stimulus. The weights of the linear mapping were selected to minimize the squared error between the decoded images and the images presented. The linear decoder was then applied to a distinct set of 150 test images. Both the predicted responses of an encoding model inferred from electrical features and the predicted responses of the fitted encoding model applied to the training images were considered. Similarity between images decoded from inferred model responses and images decoded from fitted model responses was assessed visually and by the correlation with the test images.

## Data and Code Availability

The data/code that support the findings of this study are available from the corresponding author upon reasonable request.

## Author Contributions

NPS, GG, SM, ARG, AK, NB, and EJC collected the experimental data; AS and AM. developed and supported the MEA hardware and software; MZ, GA, NPS, OKT, GG, and EJC conceived the analysis; MZ, GA, NPS, OKT, and EGW performed the analysis; MZ and EJC wrote the manuscript; all authors reviewed and edited the manuscript; and EJC supervised the project.

## Notes

### Competing Interest Statement

The authors have declared no competing interest.

